# Enlarged and highly repetitive plastome of *Lagarostrobos* and plastid phylogenomics of Podocarpaceae

**DOI:** 10.1101/392357

**Authors:** Edi Sudianto, Chung-Shien Wu, Lars Leonhard, William F. Martin, Shu-Miaw Chaw

**Author notes:** Author for correspondence: Dr. Shu-Miaw Chaw, Biodiversity Research Center, Academia Sinica, Taipei 11529, Taiwan., Prof. Dr. William Martin, Institute of Molecular Evolution, Heinrich-Heine-Universitaet Düsseldorf, Universitätsstr. 1, 40225 Duesseldorf, Germany. **Data deposition:** The complete plastid genomes of eight Podocarpaceous genera have been deposited at DDBJ under the accession numbers AP018899-AP018906. **Declarations of interest:** none.

## Abstract

Podocarpaceae is the largest family in cupressophytes (conifers II), but their plastid genomes (plastomes) are poorly studied, with plastome data currently existing for only four of the 19 Podocarpaceous genera. In this study, we assembled the plastomes from representatives of eight additional genera, including *Afrocarpus, Dacrydium, Lagarostrobos, Lepidothamnus, Microstrobos, Phyllocladus, Prumnopitys*, and *Saxegothaea*. We found that *Lagarostrobos*, a monotypic genus native to Tasmania, has the largest plastome among any cupressophytes studied to date (151,496 bp). Plastome enlargement in *Lagarostrobos* coincides with increased intergenic spacers, repeats, and duplicated genes. Among Podocarpaceae, *Lagarostrobos* has the most rearranged plastome, but its substitution rates are modest. Plastid phylogenomic analyses clarify the positions of previously conflicting Podocarpaceous genera. Tree topologies firmly support the division of Podocarpaceae into two sister clades: (1) the Prumnopityoid clade and (2) the clade containing Podocarpoid, Dacrydioid, *Microstrobos*, and *Saxegothaea*. The *Phyllocladus* is nested within the Podocarpaceae, thus familial status of the monotypic Phyllocladaceae is not supported.

## 1. Introduction

Plastid genomes (plastomes) of seed plants have an average size of 145 kb and contain highly conserved structure, including two large inverted repeats (IRs) and two single copy (SC) regions (Jansen and Ruhlman, 2012). In seed plants, the smallest plastome was found in a parasitic plant, *Pilostyles* (11 kb; Bellot and Renner, 2015), while the largest one resides in *Pelargonium transvaalense* (242 kb; Tonti-Filippini et al., 2017) with the IR longer than 70 kb. Plastid gene order is highly syntenic among seed plants (Jansen and Ruhlman, 2012), except for some lineages, such as Jasminae (Lee et al., 2007), *Trifolium subterraneum* (Cai et al., 2008), *Trachelium caeruleum* (Haberle et al., 2008), Geraniaceae (Guisinger et al., 2011), and cupressophytes (Wu and Chaw 2014, 2016; Chaw et al. 2018). In Geraniaceae, the degree of plastomic inversions is positively associated with the repeat abundance and their sizes (Weng et al., 2014). Moreover, nucleotide substitution rates and inversion frequencies are positively correlated in the plastomes of Geraniaceae (*d_N_* only; Weng et al., 2014) and cupressophytes (both *d_N_* and *d_S_;* Wu and Chaw, 2016).

Generally, plastid protein-coding genes are single copy, except those in IRs (Xiong et al., 2009). Seed plant IRs vary in size from 20 to 30 kb (Palmer, 1990), and their expansion or contraction influences the number of duplicate genes as well as plastome size. Typically, 15–17 duplicate genes are located in the IR of seed plants (Zhu et al., 2016). However, extreme IR contraction or expansion has led to only one or even >50 genes located in the IR of Pinaceae (Wakasugi et al. 1994; Lin et al., 2010; Sudianto et al., 2016) or *Pelargonium* x *hortorum* (Chumley et al., 2006), respectively. Outside the IR, plastid gene duplication has been reported in *Pinus thunbergii* (Wakasugi et al., 1994), *Trachelium caeruleum* (Haberle et al., 2008), *Euglena archaeoplastidiata* (Bennett et al., 2017), *Monsonia emarginata* (Ruhlman et al., 2017), and *Geranium phaeum* and *G. reflexum* (Park et al., 2017). Nevertheless, mechanisms underlining gene duplication in SC regions remain poorly known.

Having lost their IR (Wu et al., 2011), cupressophytes (conifers II) offer excellent resources to study the evolution of plastome size and gene duplication that are irrelevant to IR expansion or contraction. It has been revealed that in cupressophytes, synonymous substitution rates are negatively correlated with plastome size (Wu and Chaw, 2016), in agreement with the mutational burden hypothesis (Lynch, 2006). In addition, the cupressophyte plastomes are highly rearranged with the lowest GC-content among gymnosperms (Chaw et al., 2018). Only a few duplicate genes were reported in the cupressophyte plastomes, including *trnQ-UUG* (Yi et al., 2013; (Guo et al., 2014), *trnN-GUU* (Vieira et al., 2014, 2016; Wu and Chaw, 2016), *trnI-CAU* and partial *rpoC2* (Hsu et al., 2016), *trnD-GUC* (Vieira et al., 2016), and *rrn5* (Wu and Chaw, 2016). Most of these duplicate genes are involved in generating family-specific short IRs that are capable of mediating homologous recombination (Wu and Chaw, 2016).

Podocarpaceae is the largest cupressophyte family, including three major clades (Dacrydioid, Podocarpoid, and Prumnopityoid), 18–19 genera, and 173–187 species (Biffin et al. 2011; Cernusak et al., 2011; Christenhusz and Byng, 2016). Podocarpaceous species are mainly distributed in the Southern Hemisphere, with the species diversity hotspot in Malesian (Enright and Jaffré, 2011). Previous phylogenetic studies were inconclusive or controversial as to the positions of some Podocarpaceous genera, such as *Lagarostrobos, Lepidothamnus, Phyllocladus*, and *Saxegothaea* (Figure 1). For example, Conran et al. (2000) and Knopft et al. (2012) placed *Saxegothaea* as the first diverging genus of Podocarpaceae, but six other studies did not recover the same result. Moreover, *Phyllocladus* was once treated as a monogeneric family (the so-called Phyllocladaceae), separate from Podocarpaceae on the basis of morphological characteristics, chromosome numbers, and phytochemistry (Keng, 1977, 1978; Molloy, 1996; Page, 1990). Nonetheless, various molecular phylogenetic studies congruently held the genus belonging to Podocarpaceae (Conran et al., 2000; Kelch, 2002; Biffin et al., 2011). The classification of Prumnopityoid clade also varies across studies, e.g. Sinclair et al. (2002) and Biffin et al. (2011, 2012) only included *Lagarostrobos, Manoao, Parasitaxus, Halocarpus*, and *Prumnopitys* in the clade. Some authors, however, also added *Lepidothamnus* (Knopft et al., 2012) or *Phyllocladus* (Conran et al., 2000; Knopft et al. 2012) into the clade (see Figure 1).

**Figure 1.**
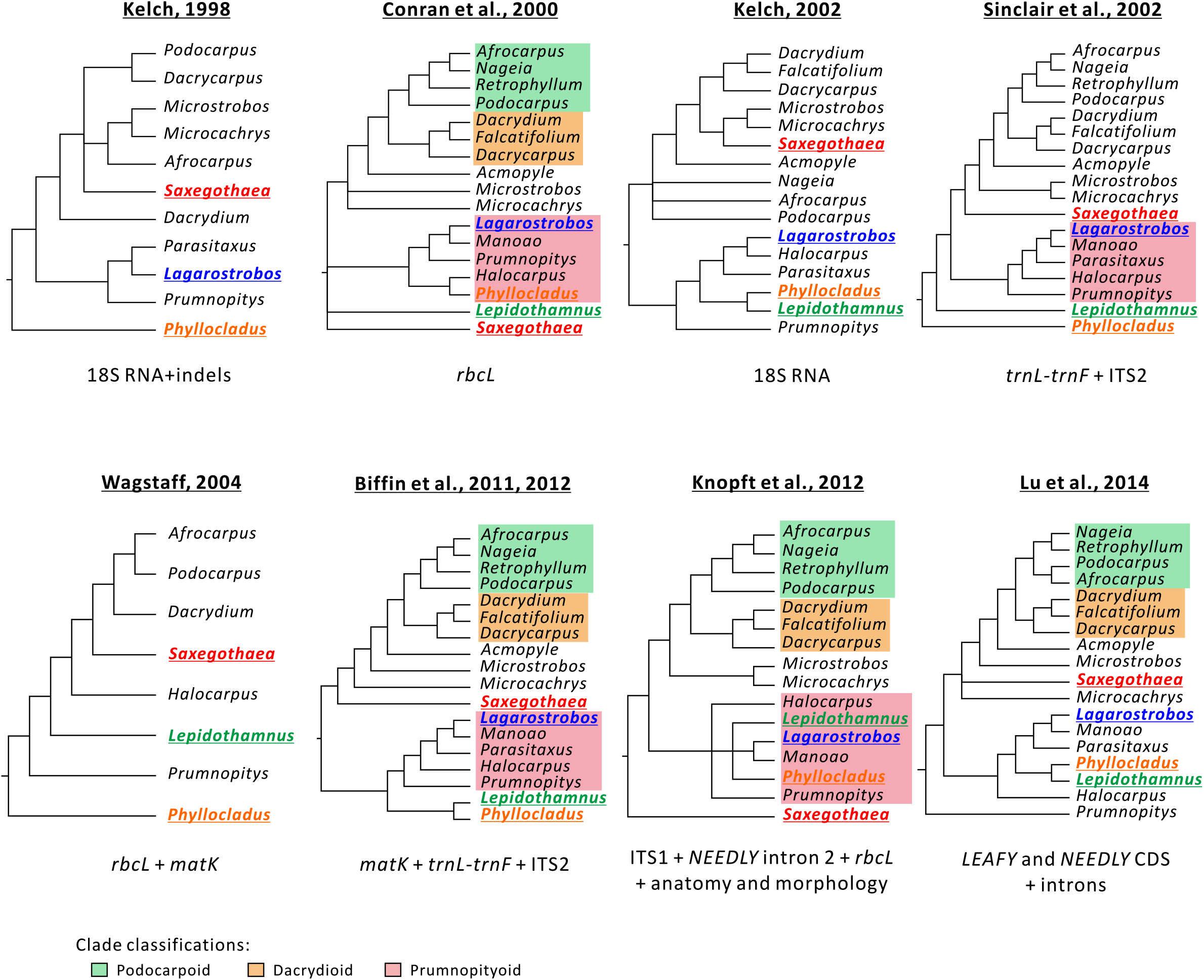
Previously reported phylogeny of Podocarpaceous genera. The eight studies have competing placements of *Lagarostrobos, Lepidothamnus, Phyllocladus*, and *Saxegothaea*.

Plastome diversity and evolution in the Podocarpaceae have yet to be systematically studied. To date, only the plastomes of the four genera – *Dacrycarpus, Nageia, Podocarpus*, and *Retrophyllum*, which are of the Dacrydioid and Podocarpoid clades, have been reported. However, samplings of the two clades are incomprehensive, and those of the Prumnopityoid clade have never been elucidated before. These may lead us to underestimate the plastome complexity in the Podocarpaceae. To this end, we have sequenced the complete plastomes from another eight Podocarpaceous genera with four representatives from the Prumnopityoid clade. The aim of this study was to (1) better sample Podocarpaceous plastome diversity and evolution, and (2) reassess intergeneric relationships across Podocarpaceae, with a particular focus on the phylogenetic position of *Phyllocladus*.

## 2. Materials and Methods

### 2.1. Plant Materials and DNA Extraction

Plant materials were collected from eight Podocarpaceous species growing in Botanischer Garten der Heinrich-Heine-Universität Düsseldorf and University of California Botanical Garden. Table S1 summarizes their collection information, voucher numbers, and GenBank accessions. For each species, total DNA was extracted from 2 grams of fresh leaves with a modified CTAB method (Stewart and Via, 1993).

### 2.2. Plastome Sequencing, Assembly, and Annotation

Sequencing tasks were conducted on an Illumina NextSeq 500 platform at Genomics BioSci & Tech (New Taipei City) or Tri-I Biotech (New Taipei City) to generate approximately 2–4 Gb of 150 bp paired-end reads for each species. The software Trimmomatic 0.36 (Bolger et al., 2014) was used to remove adapters and trim the raw sequencing reads. We used Ray 2.3.1 (Boisvert et al., 2010) for *de novo* assembly. For each species, assembled scaffolds were BLAST-searched against the *Nageia nagi* plastome (AB830885), and those with E-value < 10^−10^ were considered as plastomic scaffolds. Gaps within or between plastomic scaffolds were closed using GapFiller 1.10 (Nadalin et al., 2012) or PCR amplicons obtained from specific primers. Plastome annotations were conducted in Geneious 11.0.5 (Kearse et al., 2012) using the annotated *N. nagi* plastome as the reference, followed by manually adjusting the gene/exon boundaries. Transfer RNA (tRNA) genes were further confirmed with use of tRNAscan-SE 2.0 (Lowe and Chan, 2016). We used REPuter (Kurtz et al., 2001) to explore repetitive sequences with minimal size of 8 bp. Pseudogenes were annotated using BLASTn search against the NCBI non-redundant (nr) database.

### 2.3. Sequence Alignment and Phylogenetic Tree Reconstruction

Sequences of the 81 common plastid protein-coding genes were extracted from the eight newly sequenced and other publicly available plastomes, including four Podocarpaceous and three Araucariaceous species (Table S1). Each gene was aligned using MUSCLE (Edgar, 2004) implemented in MEGA 7.0 (Kumar et al., 2016), with the “Align Codon” option. We used SequenceMatrix (Vaidya et al., 2011) to concatenate the alignments, yielding a supermatrix that contained 84,798 characters for tree construction. Maximum likelihood (ML) and Bayesian inference (BI) trees were estimated using RAxML 8.2.10 (Stamatakis, 2014) and MrBayes 3.2.6 (Ronquist and Huelsenbeck, 2003) under a GTR + G model suggested by jModelTest 2.1.10 (Darriba et al., 2012), respectively. The bootstrap support (BS) on ML tree was assessed from 1,000 replicates. The BI analysis was run for 2,000,000 generations, in which a tree was sampled per 100 generations. The first 25% of the sampled trees were discarded as the burn-in, while the remaining ones were used to calculate the Bayesian posterior probabilities (PP).

### 2.4. Identification of Locally Collinear Blocks and Reconstruction of Ancestral Plastomes

The software progressiveMauve (Darling et al., 2010) was used to identify locally collinear blocks (LCBs) between the 12 Podocarpaceous species and *Cycas taitungensis* plastomes (AP009339). IR_A_ sequences were removed from the *Cycas* plastome prior to the LCB identification as previous studies indicated that the cupressophyte plastomes have lost IR_A_ (Wu et al., 2011; Wu and Chaw, 2014). Matrix of the LCB order and ML tree were used to reconstruct the ancestral gene order by using RINGO (Feijão and Araujo, 2016). GRIMM (Tesler, 2002) was used to estimate the plastomic inversion history.

### 2.5. Estimation of Nucleotide Substitution Rates

Both non-synonymous (*d_N_*) and synonymous (*d_S_*) substitution rates were calculated using CODEML program in PAMLX (Xu and Yang, 2013). The parameters set were runmode = 0, seqtype = 1, CodonFreq = 2, estFreq = 0, model = 1, and cleandata = 1. The ML tree shown in Figure 3 was the constraint tree.

### 2.6. Analyses of Divergence Times and Absolute Substitution Rates

The relative divergence times of the Podocarpaceous species were estimated using RelTime (Tamura et al., 2012) in MEGA 7.0. We used five estimated points from TimeTree (Hedges et al., 2015) as the calibrated ages of five specific nodes (Figure S6). Absolute *d_N_* and *d_S_* substitution rates (*R_N_* and *R_S_*) were obtained by dividing *d_N_* and *d_S_* branch lengths by the estimated time along the corresponding branches. Only branches leading to the extant species were taken into account.

## 3. Results

### 3.1. *Lagarostrobos* plastome is enlarged with abundant repeats

The eight newly sequenced plastomes are 130,343–151,496 bp long, with GC content ranging from 36.6–37.6%. They contain 81–83 protein-coding genes, 33–35 tRNAs, and four rRNAs (Table 1; Figure S1). Among them, the *Lagarostrobos* plastome is the largest. However, its gene density (0.81 genes per kb) is lower than those of other seven taxa (0.86–0.91 genes per kb). As a result, *Lagarostrobos* has the highest content of intergenic spacer (IGS) among the elucidated taxa (Table 1). *Lagarostrobos* also contains unusually abundant repeats, occupying 22.2% of its plastome (Table 1; Figure 2). By contrast, the repeat content was estimated to be 1.1–5.3% in other Podocarpaceous plastomes, suggesting that proliferation of repeats is highly active in the *Lagarostrobos* plastome. Collectively, the increased IGS and repeat contents together contribute to the enlarged plastome in *Lagarostrobos*.

**Figure 2.**
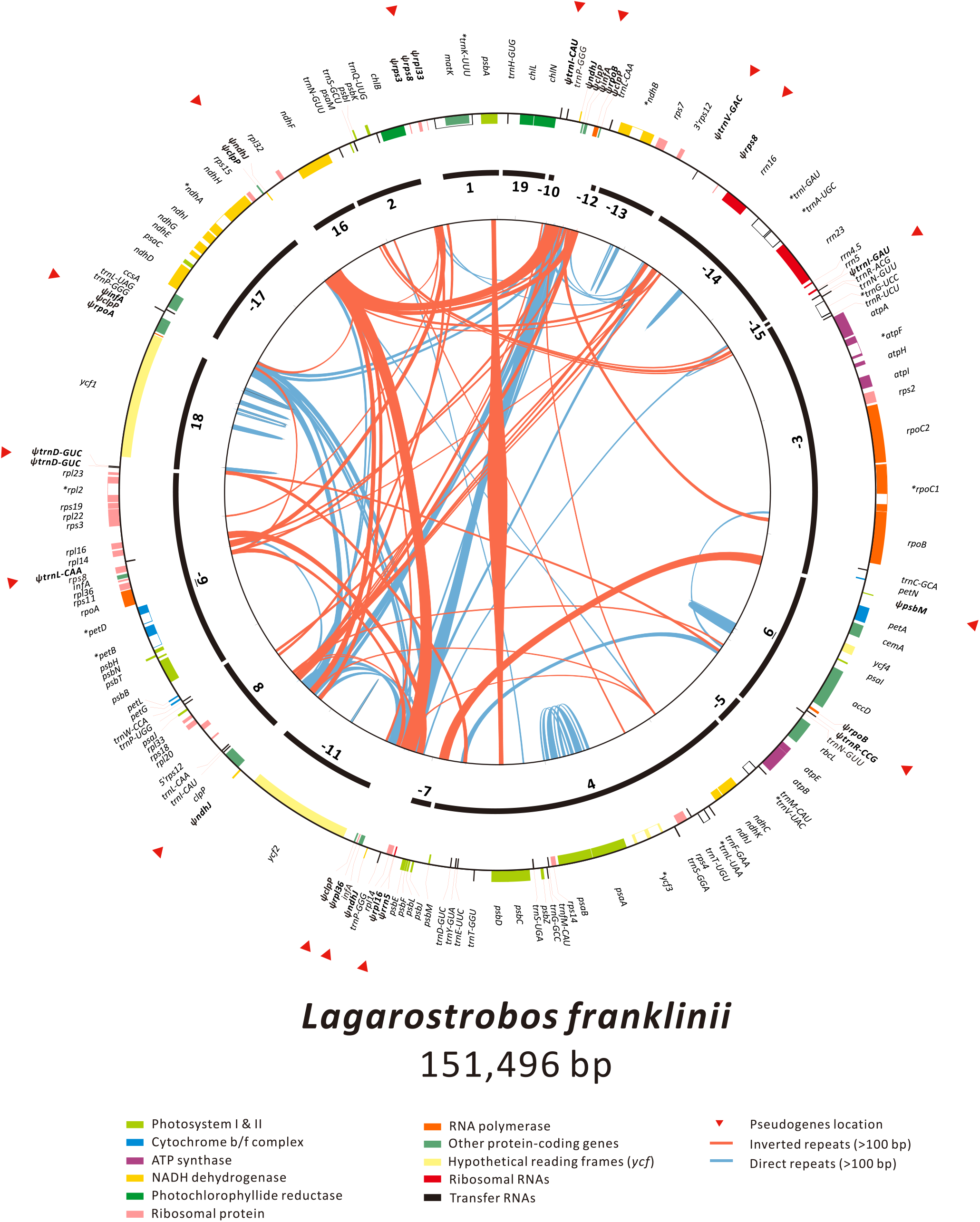
Repeat- and pseudogene-rich plastome of *Lagarostrobos Franklinii*. Colored boxes on the outermost circle represent genes being transcribed in clockwise (inner boxes) or counterclockwise (outer boxes) directions. Pseudogenes are labeled with a psi (*Ψ*). Black lines with numbers in the middle depict the locally collinear blocks (LCBs) between *Cycas* and Podocarpaceous plastomes. Negative signs indicate opposite direction as compared to *Cycas*.

**Table 1.**
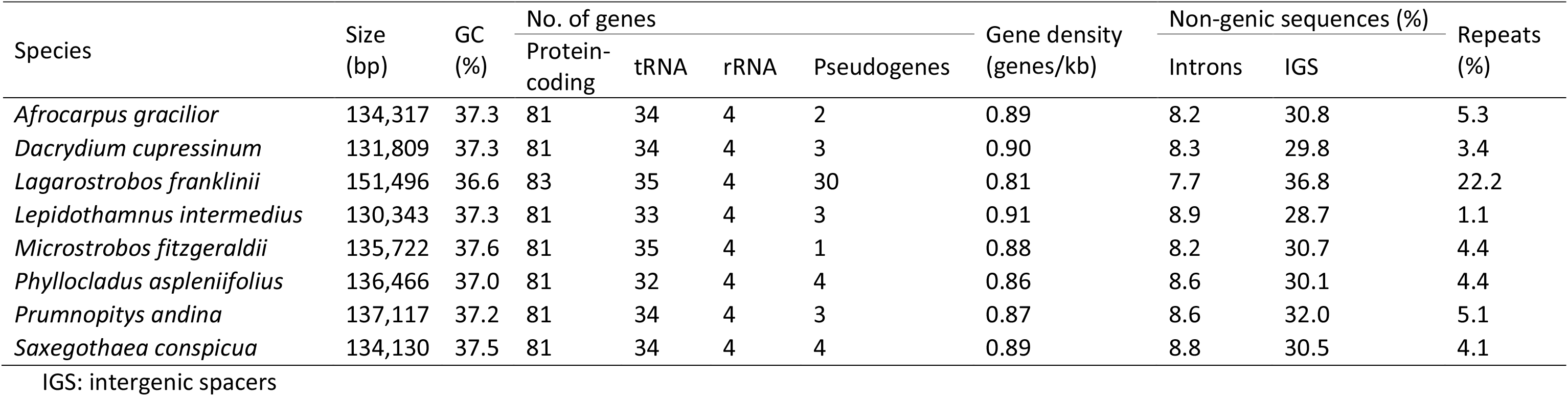
Features of the eight newly sequenced plastomes of Podocarpaceous species

A number of plastid genes vary greatly in lengths across Podocarpaceae. For example, *Lagarostrobos* and *Phyllocladus* have expanded *accD* that are 1.2 and 1.7-times longer than the average of other taxa (2,033 bp), respectively (Figure S2). In addition, the *ClpP* gene is elongated in *Lagarostrobos* and *Microstrobos* due to insertions of repetitive sequences at the 3′ end (Figure S3). Whereas, loss of introns has led to shrinkage of the *rpoC1* sequences in *Afrocarpus, Dacrycarpus, Dacrydium, Nageia, Podocarpus*, and *Retrophyllum* (Figure S1). This suggests that loss of *rpoC1* intron occurred before the divergence of the Podocarpoid and Dacrydioid clades. By contrast, *Microstrobos* is the sole taxon lacking the *atpF* intron (Figure S1).

### 3.2. Outburst of duplicate genes and pseudogenes in the Lagarostrobos plastome

Only *Lagarostrobos* has duplicated *rpl14* and *infA* (Figure 2; Figure S4). Distinct substitution rates were observed between the paralogs of these two genes, suggesting divergent evolution after the gene duplication. *Lagarostrobos* has experienced several lineage-specific duplications of plastid tRNAs, including three functional *trnP-GGG*, two functional and one pseudo (*Ψ*) *trnL-CAA*, and one functional and one *ΨtrnI-CAU* (Figure S1). In addition, *Lagarostrobos* has two *ΨtrnD-GUC* copies that are tandemly arranged.

Overall, the pseudogene content in the *Lagarostrobos* plastome amounts to ca. 3.9 kb and includes partial sequences from 22 protein-coding genes, 7 tRNAs, and 1 rRNA (Figure 2; Table 1; Table S2). Whereas, only 1–4 pseudogenes were detected in other confamilial taxa (Table 1). Most of the pseudogenes in *Lagarostrobos* are lineage-specific; only *ΨrpoB* and *ΨtrnD-GUC* are shared with other Podocarpaceous lineages (Figure S1). In *Lagarostrobos, ΨclpP* is the most copy-number abundant, with six copies scattered over the plastome (Table S2). They roughly match with three distinct regions of the functional *clpP* (Figure S5), with the mutation rates varying from 0.061 to 0.156 mutations per site.

Taken together, the *Lagarostrobos* plastome had multiple rounds of gene duplication and pseudogenization, which are unprecedented not only within Podocarpaceae but also among cupressophyte families.

### 3.3. Plastid phylogenomic analysis resolved two major clades in Podocarpaceae

Figure 3 shows ML and BI trees inferred from 81 protein-coding genes in 12 Podocarpaceous and three Araucariaceous species. These two trees are congruent in topology with 100% bootstrap support (BS) and 1.0 posterior probability (PP) in most of the nodes. Two major sister clades were resolved: (1) the Prumnopityoid clade, including *Phyllocladus, Lagarostrobos, Lepidothamnus*, and *Prumnopitys* and (2) the clade containing Podocarpoid *(Nageia, Afrocarpus, Retrophyllum*, and *Podocarpus)*,Dacrydioid (*Dacrycarpus* and *Dacrydium*), *Microstrobos*, and *Saxegothaea*. Within Prumnopityoid, *Prumnopitys* is the earliest-diverging genus, followed by *Lepidothamnus* and the clade consisting of *Lagarostrobos* and *Phyllocladus*. The sister-relationship of the latter two genera disagrees with the Keng’s views that treated *Phyllocladus* as a monogeneric family (Phyllocladaceae; Keng, 1977, 1978; also see Figure 1). *Saxegothaea* and *Microstrobos* are the successive sister taxa to the Podocarpoid-Dacrydioid clade. Within Podocarpoid, *Podocarpus* is the earliest divergent lineage, followed by *Retrophyllum* and the *Afrocarpus-Nageia* clade.

**Figure 3.**
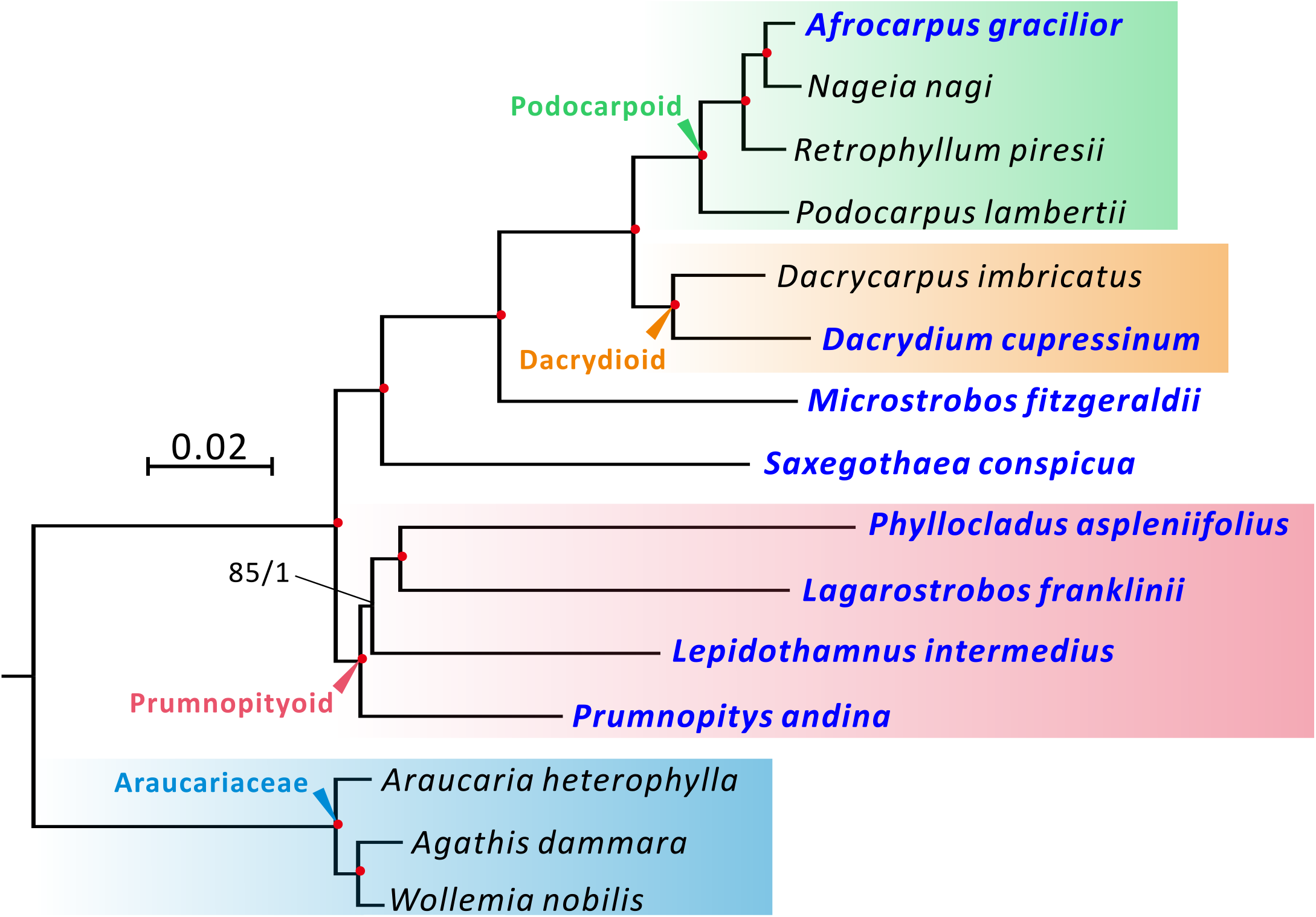
Plastid phylogenomics of Podocarpaceae. The tree framework is based on the ML tree inferred from 81 plastid protein-coding genes with three Araucariaceous species as the outgroup. Numbers along nodes indicate bootstrap support (BS)/posterior probability (PP) for ML/BI analyses. Red dots signify full support for both analyses. Newly sequenced species are highlighted in blue.

### 3.4. Podocarpaceae plastomes contain extensive inversions and diverse sets of intermediate-sized repeats

Seventeen locally collinear blocks were identified between 12 Podocarpaceous species and *Cycas* (Figure 4). The putative LCB order at each internode was inferred on the basis of the tree shown in Figure 3. Plastomic inversions that a species has experienced were estimated by comparing its LCB order with that of its ancestors. Our results show that *Lagarostrobos* has undergone at least five plastomic inversions after it diverged from *Phyllocladus* (Figure 4). Three inversions have occurred specifically in *Microstrobos*, while *Lepidothamnus* and *Podocarpus* each has undergone a lineage-specific inversion. No inversion was detected between *Saxegothaea* and the putative common ancestor of Podocarpaceae, thereby suggesting that the *Saxegothaea* plastome likely has retained the ancestral gene order in Podocarpaceae.

**Figure 4.**
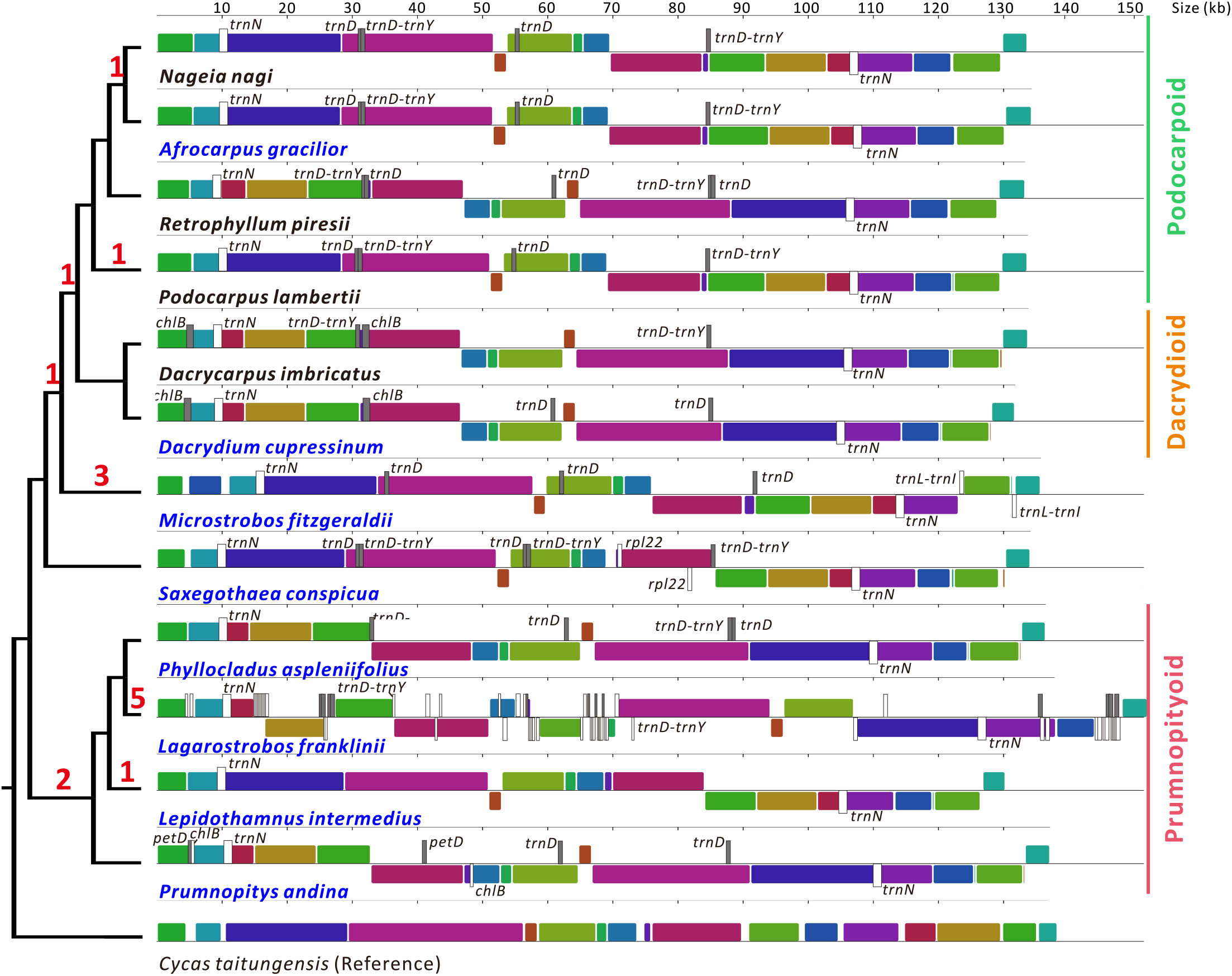
Extensive plastome rearrangements during the evolution of Podocarpaceae. The tree topology on the left side was modified from Figure 2, while colored boxes on the right side are LCBs between *Cycas* and Podocarpaceous plastomes. Boxes below horizontal lines suggest an opposing orientation relative to their counterparts in *Cycas*. Direct and inverted repeats longer than 100 bp are labeled in gray and white, respectively. Genes or gene fragments inside repeats are shown. Estimated numbers of inversion events are indicated on tree branches.

In Podocarpaceous plastomes, we found several intermediate-sized (100–1,000 bp; (Alverson et al., 2011) repeats located near the boundary of LCBs (Figure 4). All sampled taxa share an inverted repeat (IR) and a direct repeat (DR) with duplicated *trnN-GUU* (hereafter called *trnN*-IR) and *trnD-GUC* (*trnD*-DR), respectively. The *trnN*-IR and *trnD*-DR were previously reported as recombination substrates in Podocarpaceae (Vieira et al., 2016; Wu and Chaw, 2016). Here, we identify a number of novel repeats that are shared in specific clades, including *trnD-trnY*-DR in most Podocarpaceous genera, *chlB*-DR in the Dacrydioid clade, *trnL-trnI*-IR in *Microstrobos, rpl22*-IR in *Saxegothaea*, and *petD*-DR and *chlB*-IR in *Prumnopitys* (Figure 4). Overall, our data unravel a diverse set of intermediate-sized repeats in Podocarpaceae, in which *Lagarostrobos* contains the most abundant and complex repeats that are aggregated in pseudogene-rich regions (Figure 2).

### 3.5. Rearrangement frequencies do not coincide with nucleotide substitution rates in Podocarpaceae plastomes

Our dating analysis indicates that diversification of Podocarpaceae occurred around 13.39–142 MYA (Figure S6), in agreement with the viewpoint that the crown group of Podocarpaceae diversified during Mid-Jurassic to Mid-Cretaceous (Biffin et al., 2012). The absolute non-synonymous (*R_N_*) and synonymous (*R_S_*) substitution rates were estimated based on 81 concatenated genes across Podocarpaceae. In Figure 5, the estimated *R_N_* and *R_S_* rates vary from 0.13 to 0.36 substitutions per site per billion years (SSB) and from 0.43 to 0.92 SSB, respectively. *Phyllocladus* has the highest *R_N_* rate (0.36 SSB) but its *R_S_* rate (0.69 SSB) is modest, resulting in its *R_N_/R_S_* ratio (0.53) being the largest among Podocarpaceae. In contrast, *Nageia* has the fastest *R_S_* (0.92 SSB), while *Prumnopitys* has the slowest rates at both non-synonymous (*R_N_* = 0.13 SSB) and synonymous (*R_S_* = 0.43 SSB) sites. Nevertheless, none of the estimated *R_N_/R_S_* ratios of the 12 examined Podocarpaceous lineages exceeds 1, even for the highly rearranged *Lagarostrobos* (red dot in Figure 5). This finding indicates that most coding sequences of Podocarpaceous plastomes are under functional constraints, irrespective of the frequency in genome rearrangements.

**Figure 5.**
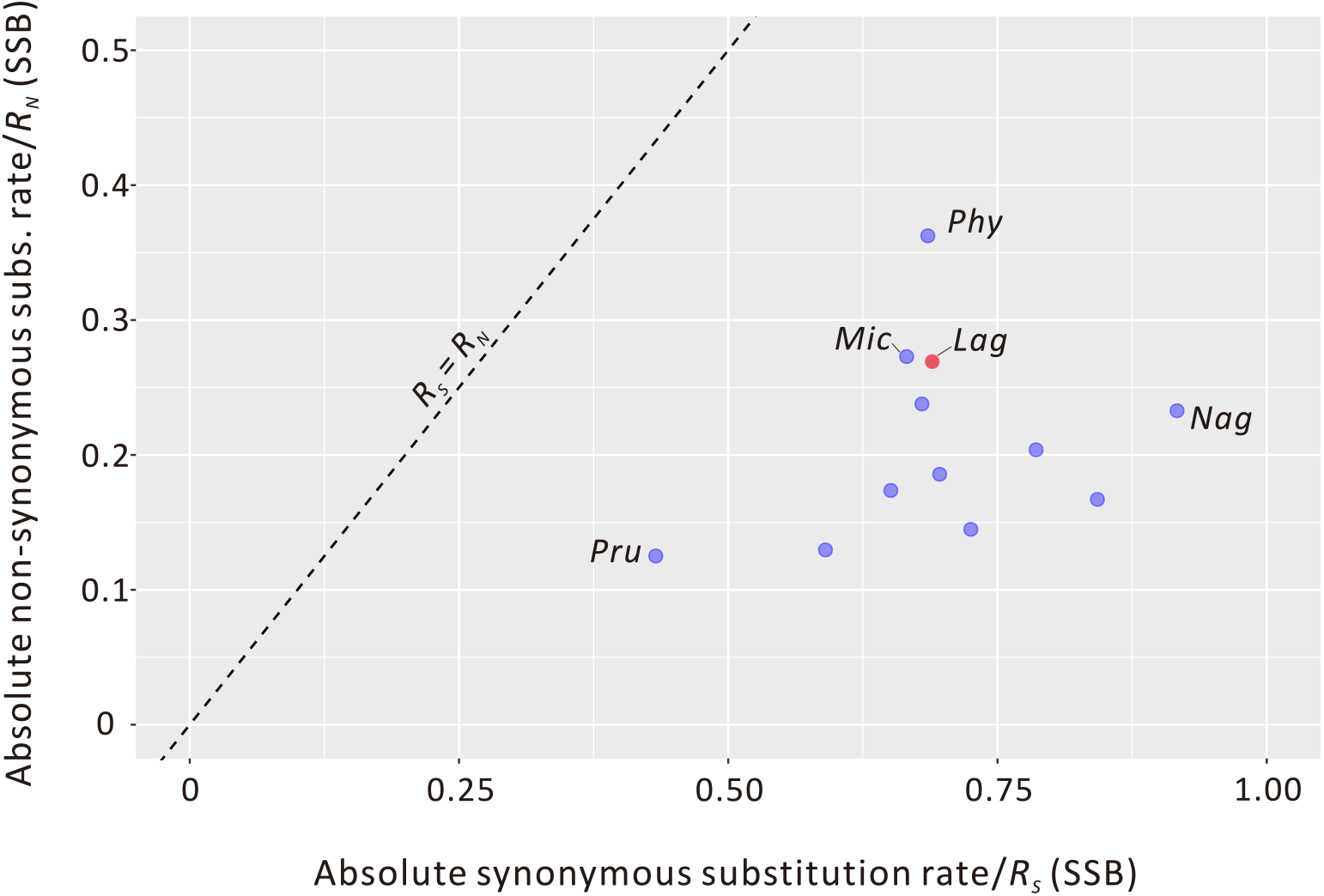
Comparison of absolute non-synonymous (R_N_) and synonymous substitution rates (Rs) estimated from plastid genes across the 12 Podocarpaceous species. The substitution rates of *Lagarostrobos* plastid genes are marked in red. SSB, substitutions per site per billion years; *Lag, Lagarostrobos Franklinii; Mic, Microstrobos Fitzgeraldii; Nag, Nageia Nagi; Phy, Phyllocladus Aspleniifolius; Pru, Prumnopitys Andina*.

## 4. Discussion

### 4.1. Insights into the phylogeny of Podocarpaceae

Most molecular phylogenetic studies of Podocarpaceae in the past were based on maximum parsimony (MP) trees inferred from a few plastid loci, e.g. *rbcL, matK, trnL-trnF*, or some nuclear genes and introns (Figure 1). But they could not agree on the inter-generic relationships in Podocarpaceae. For example, the MP trees generated by Conran et al. (2000), Sinclair et al. (2002), and Knopf et al. (2012) placed *Lepidothamnus* and/or *Saxegothaea* as the earliest divergent lineages in Podocarpaceae. This was refuted by others that placed *Phyllocladus* (Wagstaff, 2004) or *Prumnopitys* (Lu et al., 2014) as the first diverging genus. Biffin et al. 2011, 2012) divided Podocarpaceae into two major clades on the basis of three loci (Figure 1) but was still cautious about the positions of *Lepidothamnus* and *Phyllocladus* in the family.

We used 81 plastid genes to infer the phylogeny of Podocarpaceae – the largest dataset to date. Both our ML and BI trees strongly support a sister relationship between the Prumnopityoid clade and the clade containing Podocarpoid, Dacrydioid, *Microstrobos*,and *Saxegothaea* (Figure 3), and include *Prumnopitys, Lepidothamnus, Lagarostrobos*, and *Phyllocladus* in the Prumnopityoid clade – in agreement with the classification of Knopf et al. (2012), but not those of Conran et al. (2000) and Biffin et al. (2011). Therefore, our plastid phylogenomics suggests *Phyllocladus* to be a genus of Podocarpaceae rather than constituting the monotypic Phyllocladaceae. Our tree also infers that *Saxegothaea* and *Microstrobos* are the successive sisters to the Podocarpoid-Dacrydioid clade, contradicting the views of Conran et al. (2000) and Knopf et al. (2012) but agreeing with those of Sinclair et al. (2002) and Biffin et al. 2011, 2012) – see Figure 1. The intergeneric relationships within the Podocarpoid and Dacrydioid clades are highly consistent with those inferred from other studies (Conran et al., 2000; Sinclair et al., 2002; Biffin et al., 2011; Knopf et al., 2012).

### 4.2. Mechanisms underlying the enlarged Lagarostrobos plastome

We discovered that *Lagarostrobos* has a highly rearranged and enlarged plastome that contains abundant repeats (Table 1). Previously, the cupressophyte plastomes were documented to be 121.1–147.7 kb in size (Wu and Chaw 2016). With our newly sequenced data, we revised the largest of the cupressophyte plastomes to be 151.5 kb with the *Lagarostrobos* plastome holding the record.

Two mechanisms have been said to account for the enlargement of plastomes: (1) expansion of IRs (Chumley et al., 2006) and (2) accumulation of IGS sequences (Smith, 2018). We ruled out the first mechanism because *Lagarostrobos* lacks IRs (Figure 2). Instead, *Lagarostrobos* has more abundant repeats and IGS than any other Podocarpaceae (Table 1). Repeats play an important role in plastomic rearrangements (Weng et al., 2014; Sveinsson and Cronk, 2014). In addition, genomic rearrangements have been associated with increased IGS content (Sloan et al., 2012; Wu and Chaw, 2014). In *Lagarostrobos*, some long non-syntenic IGS regions (e.g. regions between LCB 11–7 and 12–10 in Figure 2) contain rich repeats, implying that they are byproducts of repeat-mediated rearrangements. We also found that, in *Lagarostrobos*, many duplicate genes are located in the repeat-rich regions (Figure 2), highlighting the key role of repeat proliferation in plastid gene duplication. Moreover, insertion of repetitive sequences accounts for the elongation of *accD* and *clpP* in *Lagarostrobos* (Figure S2, S3). As a result, repeats have led to an increase in IGS content, generation of numerous duplicate genes, and elongation of some coding genes, which together contribute to the enlarged *Lagarostrobos* plastome.

### 4.3. Abundant intermediate-sized repeats are likely responsible for numerous rearrangements in *Lagarostrobos*

Intermediate-sized repeats are able to trigger low-frequency recombination in plant mitochondrial genomes, resulting in the accumulation of substoichiometric genomes (termed sublimons), which are present at lower levels compared to the main genome (Woloszynska, 2010). The sublimons, however, may become predominant via a process known as substoichiometric shifting (SSS). Mitochondrial SSS is frequently reported in both natural and cultivated plant populations (Woloszynska, 2010). Plastomic sublimons have been reported in a number of cupressophytes, including Cupressaceae (Guo et al., 2014; Qu et al., 2017), Sciadopityaceae (Hsu et al., 2016), and Podocarpaceae (Vieira et al., 2016), and its shift is also evident among the four *Juniperus* plastomes (Guo et al., 2014). In the *Lagarostrobos* plastome, intermediate-sized repeats are far more abundant than any other Podocarpaceous genera. The abundant repeat facilitates the formation of more sublimons in *Lagarostrobos* than other genera. Notably, most of these repeats are situated at the LCB junctions, reinforcing the vital role of repeats in plastome rearrangements. Because of the presence of numerous sublimons in *Lagarostrobos*, we infer that there were at least five SSS events during the evolution of the *Lagarostrobos* plastome (Figure 4), making it as the most rearranged plastome in Podocarpaceae.

How does *Lagarostrobos* maintain such a large plastome? Smith (2016) associates bloated organellar genomes with slow mutation rates (μ) and small effective population sizes (N_e_). A recent study also indicates an inverse relationship between plastome size and *d_S_* rate in cupressophytes (Wu and Chaw, 2016). Yet, *Lagarostrobos’ R_s_* is not the lowest among Podocarpaceae (Figure 5). But as a long-lived, isolated, and highly inbred species (Shapcott, 1997), *Lagarostrobos* has an exceptionally small π (Clark and Carbone, 2008). As the substitution rates of *Lagarostrobos* plastid genes are moderate (Figure 5), we expect the small π value was mainly due to the small *N_e_*. A small N_e_ impedes the ability of natural selection to purge off excess noncoding DNA, ultimately resulting in enlarged genomes (Lynch, 2006). This viewpoint accounts for the unusually large genomes maintained in volvocine green algae (Smith et al., 2013) and similarly for the large *Lagarostrobos* plastome reported here.

## Acknowledgments

We are grateful to Dr. Sabine Etges, Botanical Garden, Heinrich-Heine-University for the kind permission in collecting samples, and to Holly Forbes and Clare W. Loughran for the help in collecting the samples in UC Botanical Garden at Berkeley. This work was supported by research grants from the ERC (666053 to WFM), the Ministry of Science and Technology Taiwan (MOST 103-2621-B-001-007-MY3 to SMC) and Biodiversity Research Center’s PI grant (2016–2018 to SMC), and the Taiwan International Graduate Program Student Fellowship (to ES).

